# Mechanisms and timing of programmed DNA elimination in songbirds

**DOI:** 10.1101/2025.05.26.656170

**Authors:** Dmitrij Dedukh, Lyubov Malinovskaya, Ondřej Kauzál, Jakub Rídl, Daria Odnoprienko, Tatyana Karamysheva, Niky Vontzou, Karel Janko, Alexander Suh, Tomáš Albrecht, Anna Torgasheva, Radka Reifová

**Author notes:** Correspondence to: Dmitrij Dedukh, Laboratory of Non-Mendelian Evolution, Institute of Animal Physiology and Genetics, Czech Academy of Sciences, Rumburska 89, 277 21 Liběchov, Czech Republic, Lyubov Malinovskaya, Department of Molecular Genetics, Cell Biology and Bioinformatics, Institute of Cytology and Genetics of the Russian Academy of Sciences, Prospekt Akademika Lavrent’yeva 10, 630090 Novosibirsk, Russia, Anna Torgasheva, Centre for Molecular Biodiversity Research, Museum Koenig Bonn, Leibniz Institute for the Analysis of Biodiversity Change, Adenauerallee 127, 53113 Bonn, Germany, Radka Reifová, Department of Zoology, Faculty of Science, Charles University, Viničná 7, 128 44 Prague 2, Czech Republic. Both authors contributed equally to the manuscript and share the first co-authorship (Dmitrij Dedukh, Lyubov Malinovskaya).

## Abstract

It is commonly assumed that multicellular organisms contain the same genetic information in all the cells of an individual. However, there is a growing list of species in which parts of the genome are removed from some cells of the organism through a process called programmed DNA elimination. In songbirds, an entire chromosome, called the germline-restricted chromosome (GRC), is lost from all somatic cells during early embryonic development. Nevertheless, the mechanisms, timing and consequences of this elimination remain largely unexplored. Here, we studied GRC elimination using two songbird species, the zebra finch (*Taeniopygia guttata*) and the Bengalese finch (*Lonchura domestica*), as model systems. We found that chromosome elimination occurs during the cleavage stage and is completed before egg laying and blastoderm formation. Elimination is associated with delayed attachment of the GRC to the mitotic spindle, changes in its histone modifications, and failure of chromatid separation in anaphase. The lagging GRC is then sequestered into a micronucleus with a defective envelope lacking the essential protein lamin B1, where the DNA is fragmented and degraded. Although the genetic basis of GRC elimination remains to be elucidated, our results suggest that changes of the GRC centromere together with epigenetic modifications of histones play a crucial role in GRC elimination from somatic cells. As the timing of elimination coincides with the germline/soma distinction, we propose that GRC elimination may play an important role in this crucial developmental process in songbirds.

## Introduction

Some organisms discard parts of their genetic material from some or all of their somatic cells during development through a process known as programmed DNA elimination (1–3). During this process, genomic regions of varying length and number are removed from the genome of somatic cells, which can lead to substantial differences in the genetic content of somatic and germ tissues (4). In some cases, such elimination concerns one or more entire chromosomes, which are removed from somatic cells and retained only in the germline. These so-called germline-restricted chromosomes (GRCs) have been observed in several animal lineages, including dipteran flies (5), hagfish and lampreys (6–8), and songbirds (9–11). Although little is known about their origin, function, or mechanisms by which they are eliminated, it has been suggested that GRCs may be involved in germline specification as well as development and that they could facilitate the resolution of germline/soma conflict (2, 5, 6, 12, 13).

Songbirds, with approximately 5,000 species, represent one of the largest known taxonomic groups where programmed DNA elimination from somatic cells occurs (14). It involves a single GRC which is eliminated not only from all somatic cells during early embryonic development, but also from male germ cells during spermatogenesis (15). For this reason, the songbird GRC is normally inherited only by females, although the occasional failure of GRC elimination in male meiosis can result in rare paternal transmission (16). The size of the GRC varies enormously, ranging from a tiny chromosome in some species to the largest chromosome in the karyotype in others (11, 14, 17). Similar to parasitic B chromosomes (18), the songbird GRC is composed of sequences that have been copied from the regular chromosomes (9, 12, 19). Although the GRC seems to be an essential part of the songbird genome for tens of millions of years, large differences in its genetic content have been observed even between closely related species, suggesting a rapid turnover of sequences on this chromosome (9, 12).

The mechanisms of GRC elimination in songbirds have so far only been studied in spermatogenesis (15), whereas embryonic tissues have not been inspected owing to methodological challenges. In male meiotic cells, the GRC is unlike regular chromosomes normally present as an univalent and is marked by histone modifications associated with heterochromatin from the early prophase of the first meiotic division, suggesting its transcriptional inactivation (20, 21). It also lacks the inner centromere protein INCENP, which plays a key role in chromosome attachment to microtubules (22). After male meiosis, the GRC can be detected as a micronucleus in the cytoplasm of secondary spermatocytes or spermatids, still retaining histone marks of condensed chromatin and showing signs of DNA fragmentation (17, 20–22). These observations suggest that GRC elimination from male germ cells may involve both epigenetic modifications and the inability of the GRC to attach to the meiotic spindle. However, the exact mechanism underlying GRC elimination from male germ cells remains unknown, and it is also unclear how and when during ontogeny the songbird GRC is eliminated from somatic cells.

In this study, we employed a combination of cytogenetic and immunohistological approaches to investigate the mechanism and timing of GRC elimination from somatic cells during embryogenesis using two songbird species, the zebra finch (*Taeniopygia guttata*) and the Bengalese finch (*Lonchura domestica*), as model systems. As part of this research, we collected embryos at multiple time points before and after oviposition, and analyzed the GRC behavior during cell divisions, along with its epigenetic modifications.

## Results

### When and how the GRC is eliminated from somatic cells

To investigate the mechanisms and timing of GRC elimination from somatic cells, we analyzed early embryos from several developmental stages in two songbird species – zebra finch (Fig. 1*A*) and Bengalese finch (Fig. 1*B*), which diverged from each other approximately 9.5 Mya (23). The analyzed embryonic stages included the cleavage stage isolated from eggs before oviposition and the blastoderm stage obtained from freshly laid eggs and eggs incubated up to 12 h after oviposition. This sampling covered embryonic developmental stages EGK.IV-IX as classified by Eyal-Giladi and Kochav (24). The GRC in embryonic cells was visualized by fluorescent *in situ* hybridization (FISH) using previously developed GRC probes for zebra finch (9, 11) and a newly developed GRC probe for Bengalese finch (*SI Appendix, Methods*).

**Figure 1.**
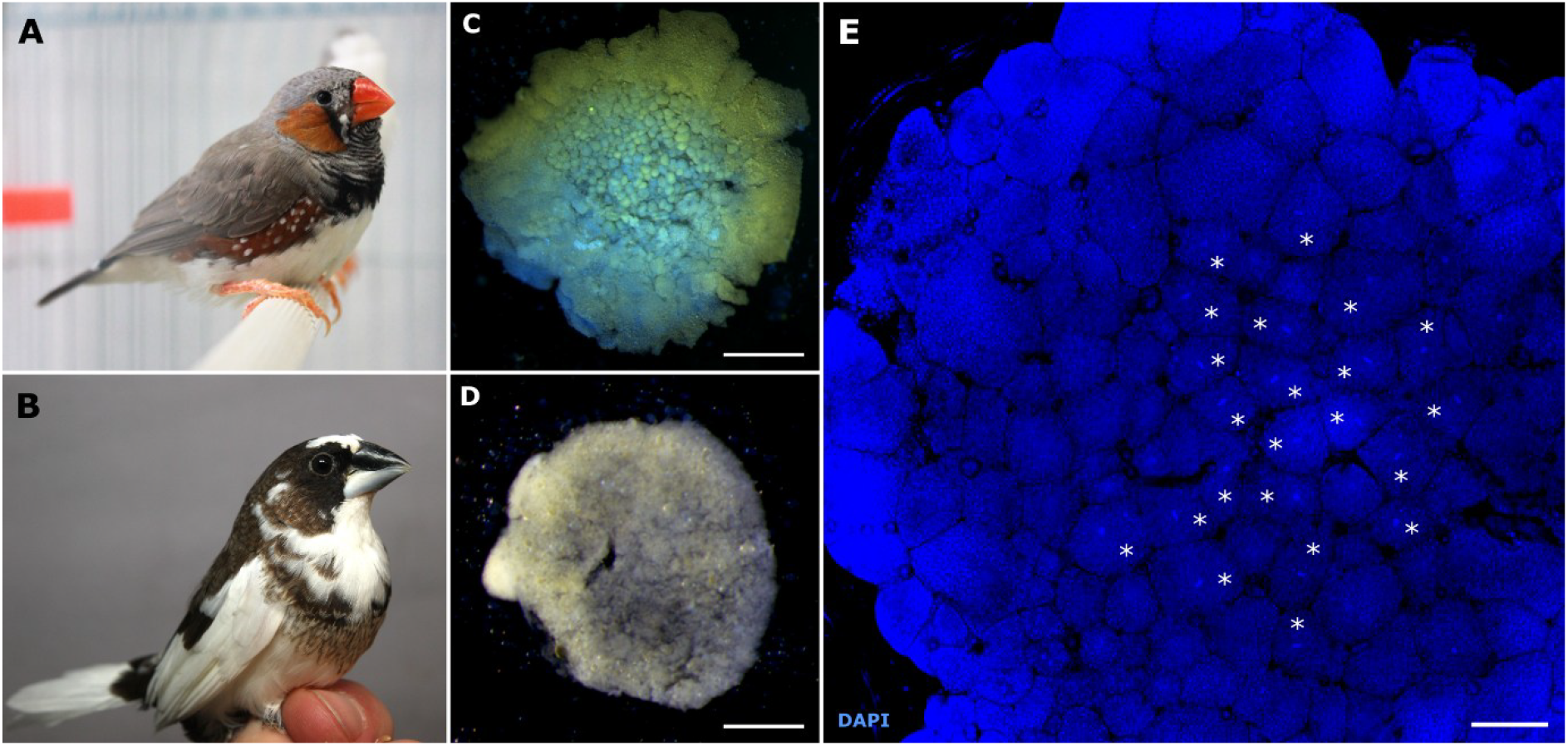
Model species, zebra finch and Bengalese finch, and their embryos at the EGK.IV and EGK.VI-IX stages. (A) Adult male zebra finch individual. (B) Adult male Bengalese finch individual. (C) Bengalese finch EGK.IV embryo isolated from eggs before oviposition. (D) Zebra finch EGK. VI-IX embryo isolated from eggs after oviposition. (E) Detail of the Bengalese finch EGK.IV embryo. Smaller mitotically dividing cells indicated by asterisks tend to be located in the center of the embryo, while larger mostly interphase cells are observed at the periphery. DNA is counterstained with DAPI (blue). Scale - 500 µm for C, D and 10 µm for E.

The early embryos isolated from eggs dissected from the female reproductive tract about 10-12 h before oviposition contained roughly between 500 and 1000 cells. Based on morphology and the number of cells, we classified these embryos as EGK.IV (Fig. 1*C*) (25). At this stage, 43%-60% of the interphase cells analyzed contained the GRC in the nucleus (*SI Appendix*, Table S1). The rest of the interphase cells were devoid of the GRC, indicating that the elimination process had already started prior to this stage. We observed that 23-40% of the cells in the visible layer of EGK.IV embryos were mitotically dividing, with most of them (75-96%) containing the GRC. Interestingly, the mitosis seemed to be synchronized among cells as in some embryos, we observed almost all cells in metaphase, while in others in anaphase. Mitotically dividing cells tended to be located in the central part of the embryos and were smaller than interphase cells located at the periphery (Fig. 1*E*).

Interestingly, we observed misalignment of the GRC relative to other chromosomes (Fig. 2*A*) in approximately one-third of dividing cells at metaphase. This may indicate a delay in GRC attachment to the mitotic spindle. During anaphase, the GRC was clearly lagging behind other chromosomes (Fig. 2*B* and *SI Appendix*, Fig. S1*A* and S1*B*). While other chromosomes were already reaching the poles of the cell, the GRC was still located in the equatorial position with its chromatids connected to each other. This suggests that failure of chromatid separation may contribute to GRC lagging. In the vast majority of cells, nuclear division and GRC lagging occurred symmetrically, apparently resulting in the loss of the GRC from both daughter cells. Only in one cell from 193 cells with GRC, both GRC chromatids seemed to segregate to the same pole, leading to duplication of the GRC in one cell and elimination in another (*SI Appendix*, Fig. S1*C*). We found no anaphase or telophase cells in EGK.IV embryos in which the GRC chromatids showed normal equal segregation together with other chromosomes. This suggests that all mitotically dividing cells eliminate (or in rare cases duplicate) the GRC at this stage.

**Figure 2.**
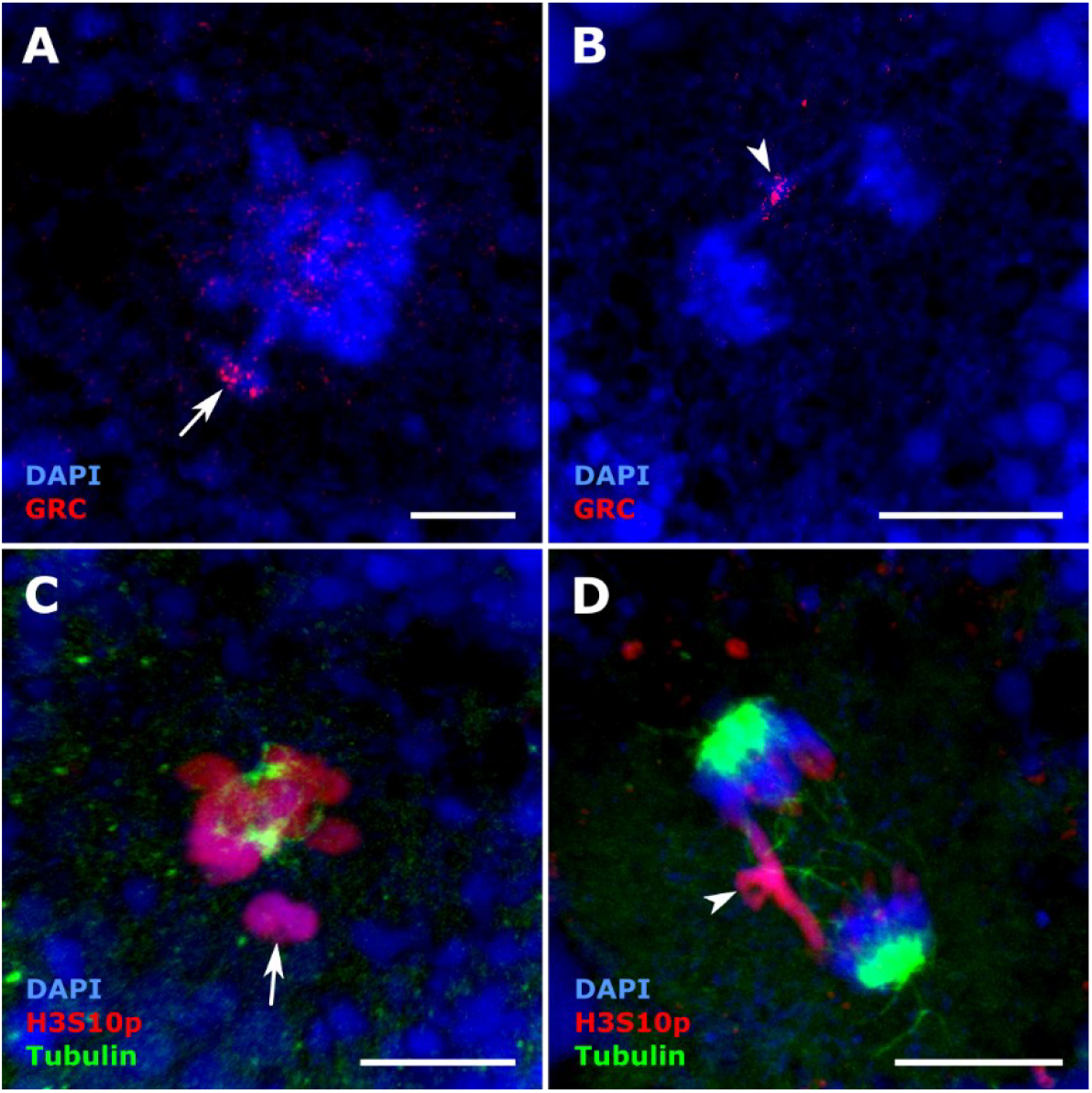
Mechanisms of GRC elimination in the zebra finch EGK.IV embryos isolated from eggs before oviposition. (A-D) Mitotically dividing chromosomes in the whole-mount embryos. The GRC is located away from the regular chromosomes at metaphase (A, C) and is lagging in anaphase (B, D). The GRC is visualized by FISH with a GRC-specific probe (red) (A, B). Embryonic cells are immunostained with antibodies against H3S10p (red) and tubulin (green) (C, D). DNA is counterstained with DAPI (blue). The arrows point to the misaligned GRCs in metaphase (A, C). Arrowheads point to the lagging GRC in anaphase (C, D). Scale - 10 µm. The same results were observed in both model species. Therefore, we only show here photos for the zebra finch. Photos from the Bengalese finch embryos can be found in the *SI Appendix*.

To investigate the epigenetic modifications of the GRC, we immunolocalized H3 histone phosphorylation at serine 10 (H3S10p), a marker of chromosome condensation, in EGK.IV embryos (Fig. 2 *C* and *D*, and *SI Appendix*, Fig. S1*D*). H3S10p normally marks chromosomes from prophase until the onset of anaphase (26), when dephosphorylation of H3S10 initiates chromatid separation and chromosome segregation in anaphase (27, 28). We observed no differences in H3S10p staining between GRC and other chromosomes during prophase and metaphase (Fig. 2*C*). However, in anaphase and telophase, the signal was absent on the segregating regular chromosomes but present on the GRC, consistent with the observed delay of the GRC segregation in anaphase and the failure of sister chromatid separation on the GRC (Fig. 2*D*, *SI Appendix*, Fig. S1*D*).

To assess the duration of GRC elimination during development, we further analyzed the embryos isolated from freshly laid eggs and those incubated for up to six hours. These embryos were at various developmental stages, ranging from EGK.VI (Fig. 1*D*), with the estimated total cell number of around 23,000, to EGK.IX. As in previous studies (29), the developmental stage of embryos from eggs incubated for a given time was not very consistent and even at the time of oviposition there was considerable variability. Since it was not possible to evaluate the developmental stage of all the embryos sampled, we combined the data from all these embryos for further analyses. The EGK.VI-IX embryos (Fig. 3*A*) contained a significantly lower percentage of interphase cells with GRC-positive nuclei (from 1% to 21% in zebra finch and from 3% to 13% in Bengalese finch) than EGK.IV embryos (*SI Appendix*, Table S1).

**Figure 3.**
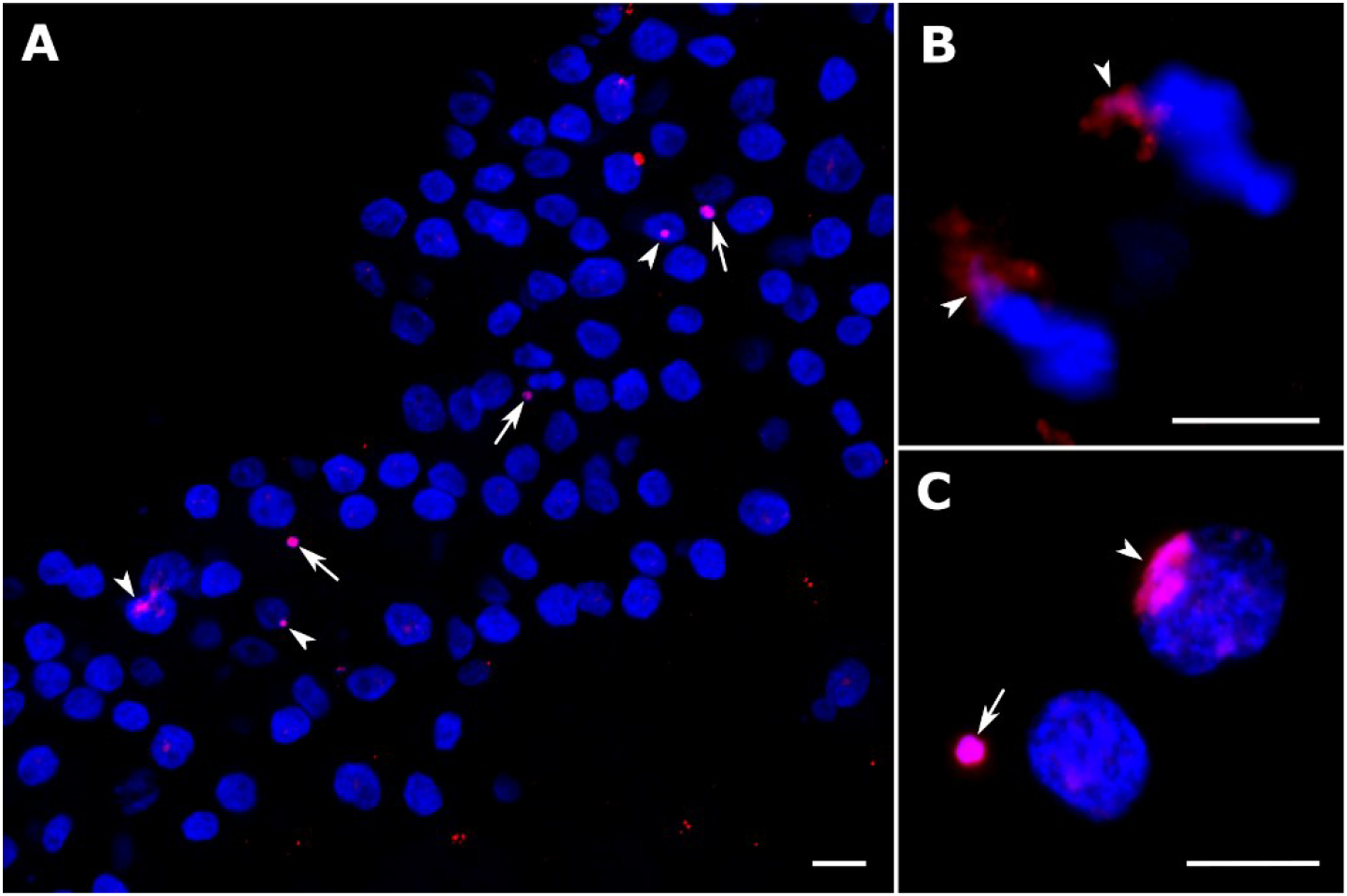
GRC in the EGK.VI-IX embryos of zebra finch, isolated from eggs after oviposition. (A) The cells with GRC-positive nuclei or micronuclei in the cytoplasm are scattered throughout the cryosectioned embryo. A photo of the entire embryo can be seen in the *SI Appendix*, Fig. S3. (B) Segregation of the GRC alongside regular chromosomes during anaphase in a whole-mount embryo. (C) Detail of the GRC in the interphase nucleus and in the micronucleus at nuclei spreads. The GRC is detected by FISH using a GRC-specific probe (red). DNA is stained with DAPI (blue). Arrowheads point to GRC-positive nuclei. Arrows point to GRC micronuclei. Scale - 10 µm.

Among the numerous mitotically dividing cells in EGK.VI-IX embryos, we did not observe a single GRC lagging in anaphase or telophase. This indicates that, by the time of oviposition, the GRC has already been eliminated from the nuclei of somatic cells and remains only in the nuclei of germ cells. In support of this, we observed the GRC segregating alongside other chromosomes in anaphase at EGK.VI-IX stage embryos (Fig. 3*B*). Cryosectioning of EGK.VI-IX embryos showed that cells with GRC-positive nuclei were scattered throughout the embryo (Fig. 3*A*, *SI Appendix*, Fig. S2), which is consistent with the previously observed distribution of primordial germ cells (PGCs) in these embryos (30). Most GRC-positive nuclei contained one GRC copy. However, two distinct GRC signals were detected in up to 7% of GRC-positive nuclei of zebra finch and Bengalese finch embryos at EGK.VI-IX stage (*SI Appendix*, Table S1 and Fig. S2).

### The fate of the GRC after its elimination from nucleus

In EGK.IV and EGK.VI-IX embryos of both species, we observed GRC-positive micronuclei (Fig. 3*A* and *C* and *SI Appendix*, Fig. S1*F-H*). This demonstrates that, after being eliminated from the nucleus, the GRC is sequestered to the micronucleus, as has been observed in spermatogenesis (10, 15). In one EGK.IV embryo, the GRC-positive micronuclei were detected in the cytoplasm of 9% of interphase cells (*SI Appendix*, Table S1). Estimates of the proportion of cells with GRC micronuclei in EGK.VI-IX embryos ranged from 3% to 7% in zebra finch and from 2% to 11% in Bengalese finch (*SI Appendix*, Table S1). We also detected some GRC-negative micronuclei, however, their proportion relative to the total number of analyzed cells was quite low (less than 1.6% in zebra finch and less than 2.4% in Bengalese finch (*SI Appendix*, Table S1). Given that our results above suggest that GRC elimination in embryogenesis is completed before oviposition, the presence of GRC-positive micronuclei in EGK.VI-IX embryos indicates that GRC micronuclei degradation requires a relatively long time.

To further trace the lifespan of GRC micronuclei, we assessed their presence and proportion at the later stages of embryogenesis, HH4 and HH8, classified by Hamburger and Hamilton (31), which were obtained from eggs incubated for 24 and 40 h, respectively. At the HH4 stage, we still detected some GRC-positive micronuclei in zebra finch (in approximately 1.8% of cells), but none in Bengalese finch. At stage HH8, no GRC-positive micronuclei were observed (*SI Appendix*, Table S1).

In male germ cells, the eliminated GRC in the micronucleus is strongly heterochromatinized (20). To test whether this also occurs in the GRC eliminated from somatic cells, we stained EGK.IV and EGK.VI-IX embryos, together with testis samples of both species, with antibodies against H3S10p and H3K9me3 histone modifications, which are associated with heterochromatin (Fig. 4 and *SI Appendix*, Fig. S3). In contrast to the GRC micronuclei in male germ cells (Fig. 4*E* and *F*, and *SI Appendix*, Fig. S3*D* and *E*), we detected only a very weak or no H3K9me3 and H3S10p signal at GRC-containing micronuclei in the embryonic cells of both zebra finch and Bengalese finch (Fig. 4*A* and *B*, and *SI Appendix*, Fig. S3*A-C*).

**Figure 4.**
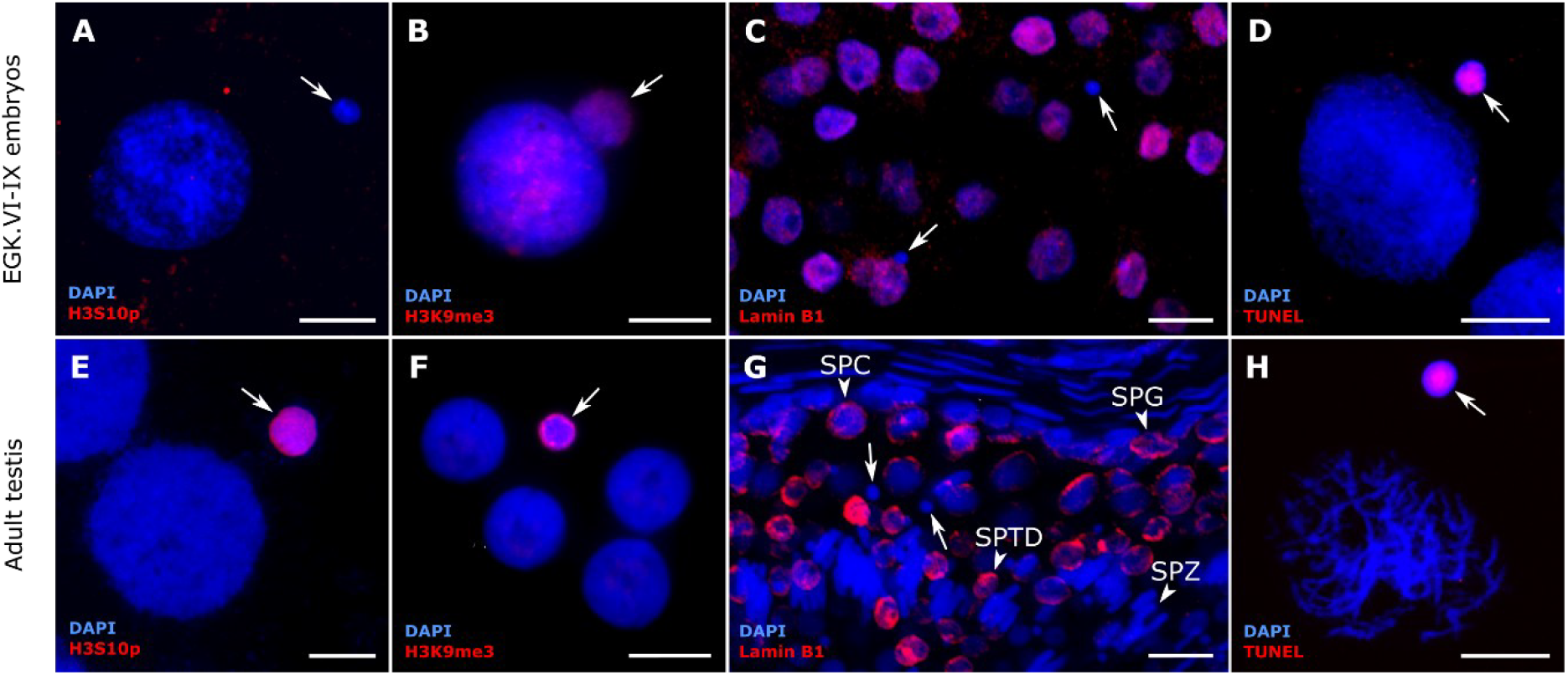
Comparison of micronuclei characteristics in EGK.VI-IX embryos and testes. (A, E) Nuclei spreads of Bengalese finch embryo and testis after immunostaining with the anti-H3S10p antibody (red). GRC micronuclei are intensively stained by anti-H3S10p antibody in testis, but not in embryos. (B, F) Nuclei spreads of zebra finch embryo and testis immunostained with the anti-H3K9me3 antibody (red). GRC micronuclei are only weakly stained in embryos, but strongly in testis. (C, G) Cryosections of zebra finch embryo and adult testis immunostained with the anti-Lamin B1 antibody (red). SPG, spermatogonium; SPC, spermatocyte; SPTD, spermatid; SPZ, spermatozoon. The antibody stains the nuclear envelope in embryonic cells as well as in spermatogonia, spermatocytes and spermatids. In contrast, the GRC micronuclei in embryos and in testis as well as nuclei of spermatozoas show no signal. (D, H) Nuclei spreads of Bengalese finch embryo and testis after TUNEL assay staining the fragmented DNA (red). The DNA is fragmented in most GRC micronuclei both in embryos and testis. DNA is stained with DAPI (blue). Arrows point to GRC micronuclei. Scale - 10 µm.

It was previously shown that micronuclei originated from various mitotic errors usually form a defective nuclear envelope lacking certain essential proteins, particularly lamin B1 (32, 33), which is also missing in the nuclear envelope in spermatids and sperm in mammals (34– 36). To check whether GRC micronuclei exhibit similar properties, we applied anti-lamin B1 antibodies on cryosections of zebra finch EGK.VI-IX embryos (Fig. 4*C*) and adult testes (Fig. 4*G*). In both cases, all micronuclei lacked the lamin B1 signal. Adult zebra finch testes also showed no lamin B1 staining of spermatozoa (Fig. 4*G*).

Finally, we examined the integrity of the DNA in the GRC micronuclei in both embryos and testes. In Bengalese finch EGK.VI-IX embryos, 76% of the micronuclei (n=41) displayed strong signal after terminal dUTP nick end labeling (TUNEL) assays (Fig. 4*D*). In testes, 68% of the micronuclei (n=122) displayed a strong positive TUNEL signal (Fig. 4*H*), indicating extensive DNA fragmentation. The rest of the micronuclei showed either weak or no signal. No DNA degradation was found in the interphase nuclei.

## Discussion

There is a growing list of species that remove parts of their genome from some cells of the organism either during embryogenesis or during gametogenesis. However, the mechanisms and evolutionary significance of such programmed DNA elimination are only beginning to be understood. Here, we investigated the mechanisms of programmed DNA elimination during the early stages of songbird embryogenesis, when the single GRC is eliminated from somatic cells. We show that GRC elimination is ongoing during cleavage in EGK.IV embryos and is completed by the time of oviposition at the EGK.VI stage. Elimination occurs during cell division, mostly in the central part of the EGK.IV embryos, and is associated with delayed attachment of the GRC to the mitotic spindle in metaphase, as well as changes in epigenetic modifications of this chromosome compared to regular chromosomes and failure of chromatid separation during anaphase. This leads to delayed segregation of the GRC to the poles of the cell and its enclosure to the micronucleus, where DNA is fragmented and degraded.

The timing as well as cytological mechanisms of GRC elimination during songbird embryogenesis are strikingly similar to those observed in other organisms. In hagfish and lamprey, GRC elimination begins at the sixth cleavage division and takes several cell divisions (37). In flies, GRCs are eliminated at the fifth cleavage division (5, 38). We have shown that GRC elimination in songbirds is ongoing in embryos containing around 500 – 1,000 cells, i.e. around the 9th and 10th cleavage divisions. However, as these embryos already contained some cells without the GRC, it is likely that GRC elimination starts already during earlier divisions. GRC elimination is completed by the time of oviposition, when the embryo contains around 23,000 cells.

Importantly, the timing of the GRC elimination in songbirds coincides with the germline/soma distinction, which is believed to occur in birds via a preformation model based on the inheritance of specific maternal factors (39). The specification of PGCs in both songbirds and chicken occurs at a similar stage of development, and the distribution of PGCs in embryos from freshly laid eggs resembles the distribution of GRC-positive cells (30, 31, 40). This also suggests that the presence of the GRC can be used as a reliable and easily detectable DNA marker of PGCs at this and subsequent stages of development. The early embryonic segregation of the immortal germline and the development of a completely disposable soma is a key innovation of animals, possibly allowing for greater evolvability of somatic tissues (39). However, we still know very little about the molecular mechanisms behind this important developmental process, especially in birds, where early embryogenesis occurs in the developing egg inside the female reproductive tract. Although the direct link still needs to be established, our results suggest that GRC elimination may play an important role in this process, likely due to the removal of certain GRC-linked genes from somatic cells. One such gene could be a paralog of the *Cpeb1* (cytoplasmic polyadenylation element binding protein 1) (12), which plays a crucial role in post-transcriptional regulation of gene expression during oogenesis and early embryogenesis, when nuclear transcription is repressed and protein synthesis largely depends on the translation of pre-stored mRNAs in the cytoplasm (41). This gene was copied to the GRC from the regular chromosomes prior to the radiation of songbirds, and the GRC copy differs from that on regular chromosomes by 86 amino-acid changes (12), suggesting that it might have evolved a new germline-specific function and regulation of a different subset of mRNAs.

Our results demonstrate that GRC elimination from somatic cells occurs during mitotic cell division and is initially associated with a delay in GRC attachment to the mitotic spindle, manifested by GRC misalignment in metaphase. The delay or inability of chromosomes to attach to the mitotic or meiotic spindle has been shown to be the underlying mechanism of chromosome elimination in several other systems, including uniparental chromosome elimination in animal and plant hybrids (4, 42, 43). The delay in GRC attachment to the mitotic spindle in songbirds might occur because the songbird GRC could contain a different centromeric sequence compared to other chromosomes, as has recently been shown in two nightingale species (44). In addition, the GRC in spermatogenesis, where elimination also occurs, has been shown to lack the inner centromere protein INCENP, which plays a pivotal role in attaching chromosome to microtubules (22). It is therefore possible that the different dynamics of kinetochore assembly, caused by different GRC centromeric sequences, may contribute to the delay in GRC attachment to the mitotic spindle.

Later in anaphase, GRC kinetochores seem to be properly attached to the microtubules and chromatids appear to be pulled in opposite directions, as indicated by the X-shaped configuration of the GRC in anaphase cells. However, the sister chromatids of the GRC show failed or delayed separation in the distal regions, suggesting that the release of sister chromatid cohesion may be impaired. This may be due to a lack of H3S10 dephosphorylation, which normally initiates chromatid separation and chromosome segregation in anaphase (27, 28). As a result, the GRC lags behind the other chromosomes during anaphase. A similar mechanism involving failure of chromatid separation and lagging in anaphase, despite presumably active centromeres, has also been observed in the case of GRC and paternal X chromosome elimination from somatic cells in the fungus gnat *Sciara coprophila* (45, 46) as well as in the case of GRCs elimination in lampreys (37). Similarly, nondisjunction of chromatids and lagging in anaphase has been shown to cause B chromosome elimination in plants (47).

The question remains as to what determines the timing of GRC elimination during ontogeny. Our observations show that virtually all mitotically dividing cells in EGK.IV embryos exhibit GRC lagging and subsequent elimination, while in later EGK.VI-IX stages, GRC segregates in mitosis along with other chromosomes. Analysis of gene expression patterns at the embryonic stages when elimination occurs could help to better understand the molecular mechanisms triggering the GRC elimination in songbirds.

The lagging GRC does not move to the poles of the dividing cells and is therefore not incorporated into new nuclei but instead forms a separate micronucleus. In contrast to spermatogenesis, where GRC micronuclei are strongly labelled by antibodies against histone modifications typical for heterochromatin, including H3K9me3 and H3S10p (17, 20–22) (Fig. 4*E* and *F*), we observed only a weak or no signal at GRC micronuclei in embryos (comparable to the regular pericentromeric staining in interphase nuclei). This suggests that the level of heterochromatinization and histone modifications in GRC micronuclei may differ between testes and early embryos. Importantly, we have shown that the DNA in GRC micronuclei in embryos is positively stained by TUNEL, demonstrating that the GRC in the micronucleus becomes fragmented and degraded, similar to what has been previously observed in GRC micronuclei in testis (22) (Fig. 4*D* and *H*).

We also showed that GRC micronuclei in both embryos and testes form a defective envelope lacking the essential protein lamin B1. This is consistent with previous findings in the sea lamprey, in which lamin B1 is also absent from GRC micronuclei (37). Defective envelope assembly lacking lamin B1 and other key nuclear envelope proteins has also been observed in micronuclei resulting from aberrant mitotic events, such as those occurring during cancer cell divisions (33, 48). These defective envelopes prevent the import of key proteins, leading to DNA damage, spontaneous envelope rupture and the subsequent micronuclei degradation (32). In cancer cells, fragmented and rearranged chromosomes in micronuclei occasionally fuse with the nucleus, leading to structural rearrangements of chromosomes (chromothripsis) (33). Interestingly, the GRC also shows very rapid evolution associated with frequent additions, duplications and deletions on this chromosome (9, 12), which could theoretically be caused by occasional reincorporation of restructured chromosomes in micronuclei back into the nucleus as was shown for some highly rearranged chromosomes in plant hybrids (49). We have shown that lamin B1 is also absent from the nuclear membrane of songbird spermatozoa, which is consistent with observations in mammals showing that this protein is progressively removed from spermatids and spermatozoa during nuclear envelope remodeling and head shaping (35, 36).

In conclusion, our results suggest that GRC elimination from somatic cells in songbirds involves multiple mechanisms, including a delay in GRC attachment to the mitotic spindle and chromosome lagging at anaphase associated with a failure of chromatid separation. Similar mechanisms of chromosome elimination have been observed in other unrelated organisms, suggesting that centromere modifications and the manipulation of chromosomes during anaphase arose independently several times during animal and plant evolution, and may represent a universal mechanism of chromosome elimination. Notably, we have shown that the timing of GRC elimination during embryogenesis coincides with the timing of germline/soma distinction, suggesting that GRC elimination may play a crucial role in this fundamental developmental process, which is particularly poorly understood in birds. Finally, we proposed that the mechanism of GRC elimination, involving the formation of micronuclei with defective envelopes, may contribute to the previously observed rapid evolution of this peculiar chromosome.

## Materials and Methods

### Experimental animals and embryo collection

Zebra finches and Bengalese finches were housed at 22-25°C with a 14h:10h day-night cycle. Breeding pairs were kept in separate cages provided with a nestling box and ad libitum hay, food and water. The care and experimental use of zebra finches were approved by the Animal Care and Use Committee of the Institute of Cytology and Genetics SB RAS (protocol # 199). The experimental use of Bengalese finches and zebra finches in Studenec were approved by the Ministry of Agriculture of the Czech Republic (approval no. MZE-50144/2022-13143).

To obtain embryos from eggs before oviposition, one zebra finch and three Bengalese finch females were sacrificed by cervical dislocation and eggs from their reproductive tract were dissected at about 18:30 in the evening, i.e. approximately 10-12 h before egg laying. According to morphology, these embryos corresponded to the EGK.IV stage. In addition, we analyzed embryos from freshly laid eggs collected once a day at the onset of the light cycle. The collected eggs were immediately processed or put in 4°C and later incubated at 37-38°C (80-95% humidity) for up to 6 hours to start the embryonic development. In agreement with previous observations (30), embryos from these eggs corresponded to EGK.VI to IX stages. We also analyzed embryos from eggs incubated for ∼24 hours, which corresponded to HH4 stage and 40 hours, which corresponded to HH8 stage. All analyzed embryos and their characteristics are listed in *SI Appendix*, Table S1.

To compare mechanism of GRC elimination in early embryos with those of in male germ cells, two reproductively active males of zebra finch and one male of Bengalese finch were sacrificed by cervical dislocation and testes were dissected for the preparation of meiotic spreads and cryosections. Additionally, one Bengalese finch male was used for sequencing of the germline (testis) and somatic (liver) genomes to develop the GRC-specific probe (see *SI Appendix*, Supplementary Methods).

### Embryo dissection

The yolks with the embryos were dissected from eggs and transferred into the PBS solution. To extract embryos, we used a piece of filter paper with a central hole following the protocol described by Ferran *et al*. (50). To remove the yolk residuals, the embryos were washed in 0.1M sucrose or PBS solution. Extracted embryos were photographed to determine their stage of development. Early stages of embryonic development were classified following the Eyal-Giladi and Kochav (EGK) system (51). To classify the embryos of the streak stages and onward we used the Hamburger and Hamilton (HH) system (52) adjusted for zebra finch (53).

### Nuclei spreading

Zebra finch and Bengalese finch embryos isolated from eggs after oviposition (stages EGK.VI-IX) (n=21) and testes from adult males (n=3) were used for nuclei spread preparations (*SI Appendix*, Table S1). The suspension of embryonic or testis cells was prepared by a micropipetting and proceeded to a drying-down method described by Peters *et al*. (54) with a few modifications. Briefly, dry clean slides were dipped into 1% paraformaldehyde (PFA) with 0.15% Triton X-100 in PBS. Around 20-30μl of cell suspension was slowly dispersed on the slide and left in a humid chamber at RT for 1-2h or at +4°C overnight. The slides were then washed twice in 0.4% Kodak Photo-flo and air-dried. Bengalese finch embryos were gently resuspended in hypotonic solution 0.075M KCl followed by fixation in Carnoy solution (3 volumes of methanol and 1 volume of acetic acid). After three fixative changes cell suspension was dropped on slides. Slides were kept at +4°C for up to a month or -20°C for longer storage (up to 1 year). To thaw the slides we left them at +4°C for 15 min, then at RT for 15 min, and then proceeded to immunostaining or FISH.

### Whole-mount preparation and histological sectioning

Intact zebra finch and Bengalese finch embryos isolated from eggs before oviposition (stage EGK.IV) (n=4) and after oviposition (stages EGK.VI-IX) (n=5) were used for whole mount fixation (*SI Appendix*, Table S1). Embryos were fixed overnight with 2% paraformaldehyde in PBS, then transferred to PBS and kept until immunostaining and FISH.

Embedding and cryosectioning (thickness, 18-20 μm) of embryos (n=5, stages from EGK.VI to HH8) and adult testis (n=1) were performed following the protocol outlined by Giandomenico *et al*. (55). The embryos were cut transversely for immunostaining and FISH analysis or longitudinally for the estimation of the total number of cells in the EGK.VI embryo.

### Preparation of GRC-specific probes, FISH, immunostaining and TUNEL assay

To label zebra finch GRC, we used the whole-chromosome DNA probe derived from microdissected GRC micronuclei (11) and FISH probe targeting previously described repeat specific to zebra finch GRC (9). To label Bengalese finch GRC, we developed FISH probe targeting tandem repeat specific to Bengalese finch GRC (see *SI Appendix*, Supplementary Methods for details). FISH was performed according to standard protocols for nuclei spreads (60), whole mount (57, 58) and cryosections (61).

Nuclei spreads were immunostained following the protocol described by Anderson *et al*. (56). Whole-mount immunostaining was performed according to the protocol described in Dedukh *et al*. (57, 58). Immunostaining of cryosections was performed using a standard procedure described by Eberhart *et al*. (59). Antibodies used in this study are listed in *SI Appendix*, Table S2.

TUNEL assay was performed on nuclei spreads of Bengalese finch EGK.VI-IX embryos and adult male testes according to the manufacturer’s protocol (ab66110, Abcam).

After FISH, immunostaining and TUNEL assay, slides were mounted with a Vectashield antifade mounting medium with DAPI (Vector Laboratories, Burlingame, CA, USA) to inhibit fluorescence fading.

### Microscopic analysis

Zebra finch and Bengalese finch nuclei spreads we analyzed using the Axioplan 2 microscope (Carl Zeiss, Jena, Germany) equipped with CHROMA filter sets, a CCD camera (CV M300, JAI Corporation, Yokohama, Japan), and the ISIS4 image-processing package (MetaSystems GmbH, Altlußheim, Germany) and using Provis AX70 microscope (Olympus, Japan) equipped with standard fluorescence filter sets, CCD camera (DP30W, Olympus, Japan), and the Olympus Acquisition Software.

To visualize cryosections we used the Olympus Fluoview FV3000 confocal laser scanning microscope and Olympus FV31S-SW viewer software. Whole-mount embryos were analyzed by confocal laser scanning microscope Leica TCS SP5 with an inverted Leica DMI 6000 CS base (Leica Microsystems, Germany). Diode, argon, and helium-neon lasers were employed to excite the fluorescent dyes DAPI, and the fluorochromes Alexa 488, and Alexa 594/Rhodamine respectively. Samples were analyzed with an HC PL APO 63× objective. Images were captured and processed using LAS AF software (version 4.0.11706, Leica Microsystems, Germany). To analyze confocal images of cryosections we used an image processing package Fiji (Fiji Is Just ImageJ) (62).

## Acknowledgements

We are grateful to Pavel Borodin for insightful comments and valuable suggestions that contributed to the development of this work. We thank members of the GRC Brainstorming group for helpful discussions. This research was funded by the Czech Science Foundation (grant 23-07287S to R.R., D.D. and T.A.), the Russian Science Foundation (grant 23-14-00182 to L.M. and D.O.). The Institute of Animal Physiology and Genetics receives support from Institutional Research Concept, Grant/ Award Number: RVO67985904. A.S. and A.T. were supported by the Bonn Institute for Organismic Biology - Animal Diversity (University of Bonn). The immunofluorescent imaging was performed using resources of the Common Facilities Center of Microscopic Analysis of Biological Objects, ICG SB RAS supported by the Institute of Cytology and Genetics (FWNR-2022-0015).

## Data availability

The novel whole genome sequencing data (10x Genomics linked-read libraries sequenced by Illumina) generated in this study have been deposited in the NCBI’s SRA database under the BioProject accession number PRJNA1248916.

## Notes

### Competing Interest Statement

The authors have declared no competing interest.

