## Supporting Information for "Mechanisms and timing of programmed DNA elimination in songbirds"

**This PDF file includes:**

Figures S1 to S3
Tables S1 to S2
Supplementary Methods
Figure S4

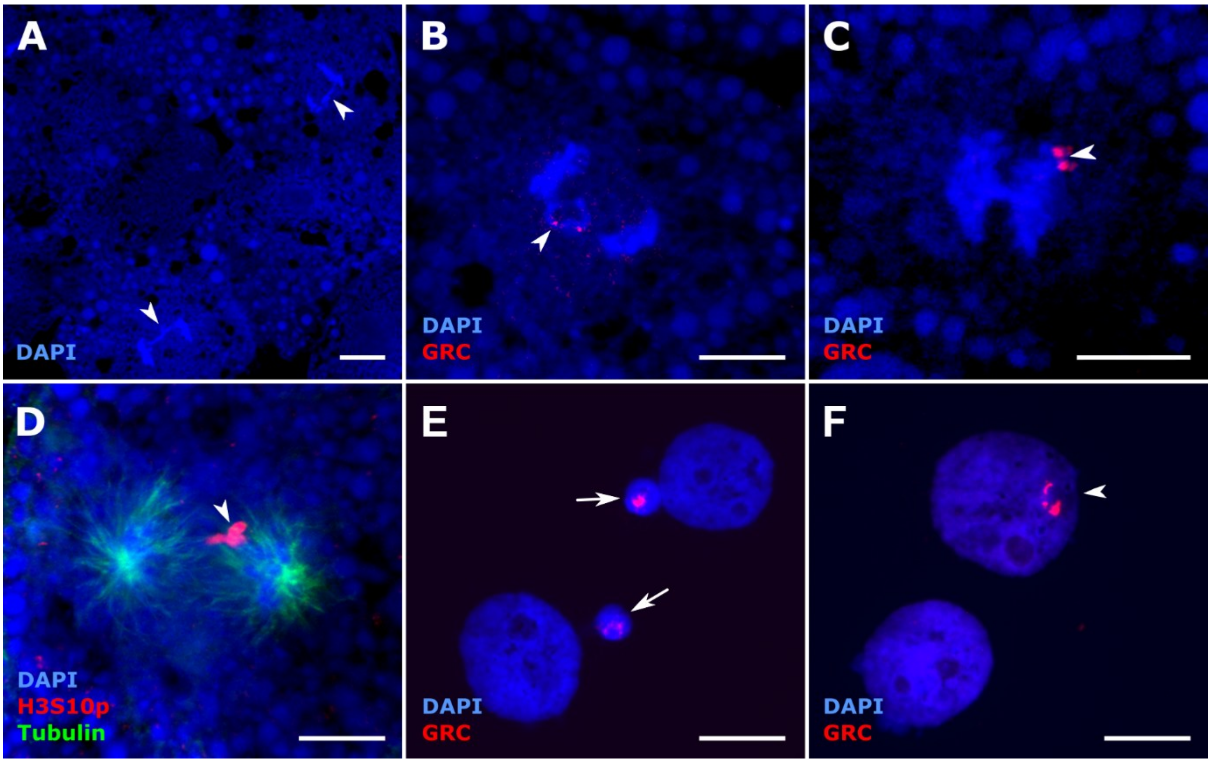


**Fig. S1.** GRC behavior in the Bengalese finch embryos. (A) Whole-mount EGK.IV embryo with anaphase cells showing a single lagging chromosome (putative GRC). (B) In the vast majority of anaphase cell of the EGK.IV embryo the lagging chromosome was positively stained with the GRC-specific probe after the FISH (red). (C) In one anaphase cell of EGK.IV embryo, we observed two GRC signals after FISH (red) at one of the cells and no signal at another pole, suggesting that in rare cases both GRC chromatids may segregate to one daughter cell. (D) The anaphase cell of the EGK.IV embryo after immunostaining with antibody against H3S10p (red) and tubulin (green). Unlike regular chromosomes, the lagging GRC is positively stained with the H3S10p antibody. (E, F) Nuclei spreads of EGK.VI-IX embryos after FISH with the GRC-specific probe (red) showing GRC positive micronuclei or nuclei. Arrowheads point to lagging GRCs (A, B, D), Two GRC chromatids segregating to one pole of the cell (C), and GRC in the interphase nuclei (F). Arrows point to GRC micronuclei (E). DNA is stained with DAPI (blue). Scale - 10 μm.

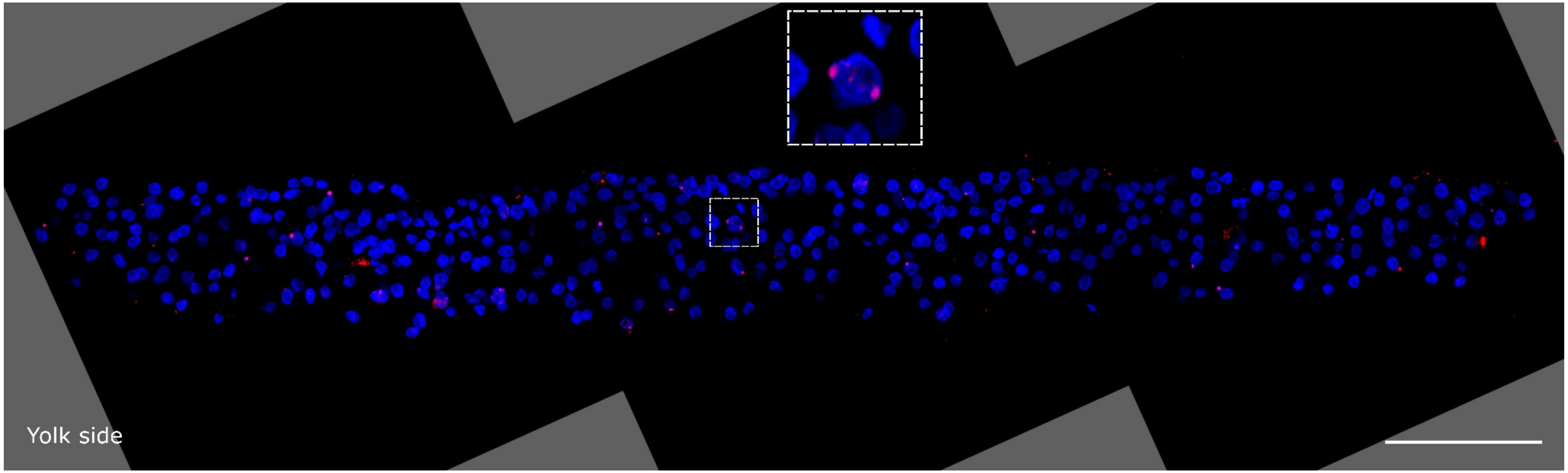


**Fig. S2.** Distribution of GRC-positive nuclei and GRC micronuclei in the cryosection of zebra finch EGK.VI-IX embryo (merge of three images).
GRC is stained by FISH with the GRC-specific probe (red). DNA is stained with DAPI (blue). Insert shows a rare cell with two GRC signals. Scale - 100 μm.

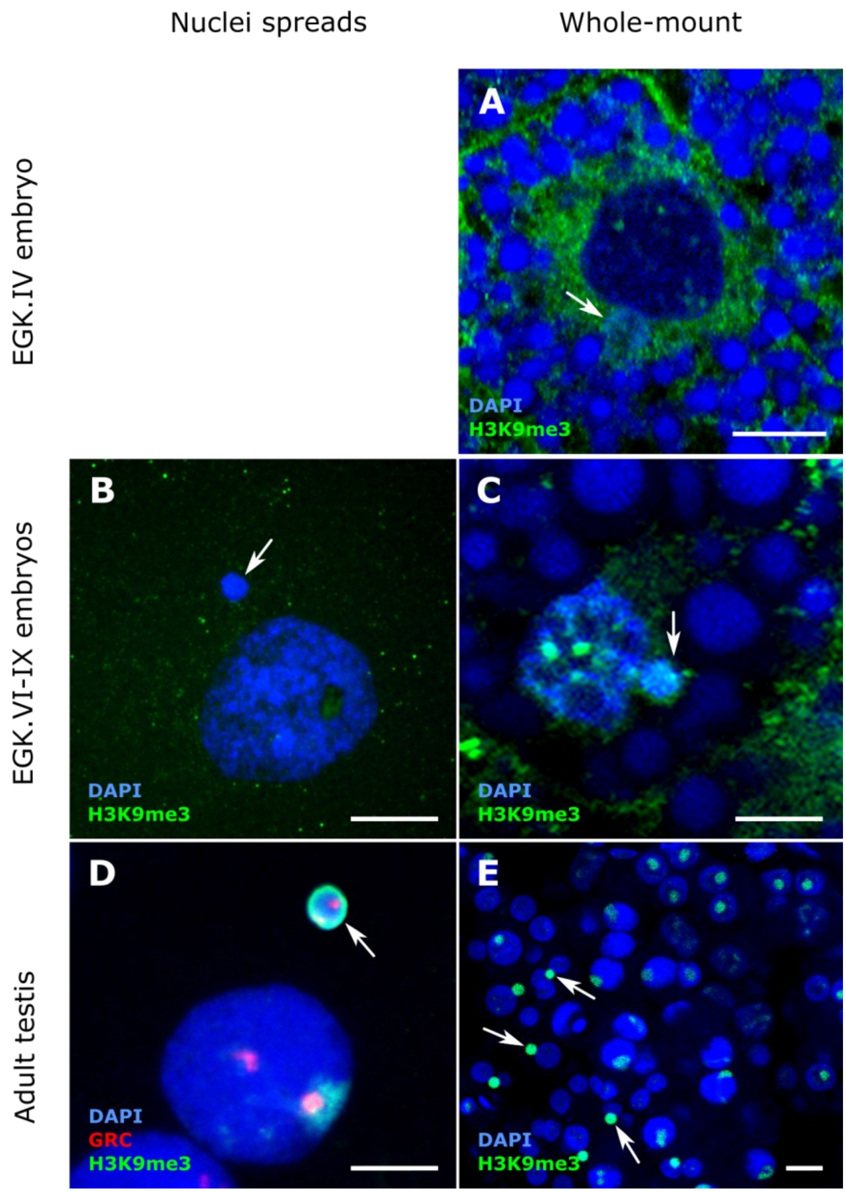


**Fig. S3.** Comparison of H3K9me3 labelling in Bengalese finch embryos and adult testes in whole-mount preparations and nuclei spreads. Samples were immunostained using the anti-H3K9me3 antibody (green) and in (D) the GRC is labelled by FISH using a GRC-specific probe (red). The GRC micronuclei showed intensive H3K9me3 signals in testes (D, E), but not in embryos (A, B, C). DNA is stained with DAPI (blue). Arrows point to GRC micronuclei. Scale – 10 µm.

**Table S1.** The list and characteristics of analyzed embryos.

| Species[1] | Stage of the embryo | Slide preparation type[2] | Sample ID | Number of cells/nuclei analyzed | Number (and proportion) of GRC-positive nuclei | Number of nuclei with 2 GRC signals[3] | Number of GRC-positive micronuclei and their proportion relative to the number of cells/nuclei[4] | Number of GRC-negative micronuclei and their proportion relative to the number of cells/nuclei[4] | Number of cells with micronuclei (without GRC staining) |
|---|---|---|---|---|---|---|---|---|---|
| ZF | EGK.IV | WM | zf_1 | 76 mitotic | 65 (85.5%) | 0 | n/a* | n/a* | |
| | | | | 111 interphase | n/a* | n/a* | n/a* | n/a* | 21 |
| BF | EGK.IV | WM | bf_1 | 72 mitotic | 60 (83.3%) | 0 | n/a** | n/a** | |
| | | | | 134 interphase | 57 (42.5%) | 0 | n/a** | n/a** | |
| BF | EGK.IV | WM | bf_2 | 28 mitotic | 21 (75.0%) | 0 | n/a** | n/a** | |
| | | | | 51 interphase | 26 (50.9%) | 0 | n/a** | n/a** | |
| BF | EGK.IV | WM | bf_3 | 49 mitotic | 47 (95.9%) | 0 | 0 | 0 | |
| | | | | 207 interphase | 125 (60.4%) | 13 | 18 (8.7%) | 0 | |
| ZF | EGK.VI-IX | NS | zf_2 | 425 | 17 (4.0%) | 0 | 29 (6.8%) | 1 (0.2%) | |
| ZF | EGK.VI-IX | NS | zf_3 | 444 | 90 (20.3%) | 0 | 13 (2.9%) | 1 (0.2%) | |
| ZF | EGK.VI-IX | NS | zf_4 | 519 | 87 (16.8%) | 0 | 27 (5.2%) | 5 (1.0%) | |
| ZF | EGK.VI-IX | NS | zf_5 | 970 | 80 (8.3%) | 0 | 72 (7.4%) | 16 (1.6%) | |
| ZF | EGK.VI-IX | NS | zf_6 | 1028 | 218 (21.2%) | 8 | 72 (7.0%) | 9 (0.9%) | |
| ZF | EGK.VI-IX | NS | zf_7 | 814 | 90 (11.1%) | 6 | 34 (4.2%) | 5 (0.6%) | |
| ZF | EGK.VI-IX | NS | zf_8 | 619 | 85 (13.7%) | 5 | 28 (4.5%) | 5 (0.8%) | |
| ZF | EGK.VI-IX | NS | zf_9 | 847 | 18 (2.1%) | 0 | 40 (4.7%) | 7 (0.8%) | |
| ZF | EGK.VI-IX | NS | zf_10 | 856 | 8 (0.9%) | 0 | 52 (6.1%) | 7 (0.8%) | |
| ZF | EGK.VI-IX | TC | zf_11 | 4882 | 85 (1.7%) | n/a | 261 (5.3%) | 43 (0.9%) | |
| ZF | EGK.VI-IX | TC | zf_12 | 4252 | 43 (1.0%) | n/a | 163 (3.8%) | 20 (0.5%) | |
| ZF | EGK.VI-IX | LC | zf_13 | 23141 | n/a | n/a | n/a | n/a | |
| ZF | EGK.VI-IX | WM | zf_14 | n/a | n/a* | n/a* | n/a* | n/a* | |
| BF | EGK.VI-IX | NS | bf_4 | 1172 | 77 (6.6%) | 0 | 78 (6.7%) | 11 (0.9%) | |
| BF | EGK.VI-IX | NS | bf_5 | 1049 | 134 (12.8%) | 5 | 86 (8.1%) | 10 (1.0%) | |
| BF | EGK.VI-IX | NS | bf_6 | 1199 | 141 (11.8%) | 2 | 67 (5.6%) | 12 (1.0%) | |
| BF | EGK.VI-IX | NS | bf_7 | 219 | 14 (6.3%) | 0 | 17 (7.8%) | 1 (0.5%) | |

| | | | | | | | | | |
|---|---|---|---|---|---|---|---|---|---|
| BF | EGK.VI-IX | NS | bf_8 | 974 | 48 (4.9%) | 1 | 71 (7.3%) | 8 (0.8%) | |
| BF | EGK.VI-IX | NS | bf_9 | 1622 | 51 (3.1%) | 3 | 126 (7.8%) | 20 (1.2%) | |
| BF | EGK.VI-IX | NS | bf_10 | 2284 | 79 (3.5%) | 4 | 122 (5.3%) | 29 (1.3%) | |
| BF | EGK.VI-IX | NS | bf_11 | 645 | 43 (6.7%) | 0 | 48 (7.4%) | 2 (0.3%) | |
| BF | EGK.VI-IX | NS | bf_12 | 770 | 36 (4.7%) | 1 | 18 (2.3%) | 3 (0.4%) | |
| BF | EGK.VI-IX | NS | bf_13 | 112 | 4 (3.5%) | 0 | 7 (6.3%) | 0 | |
| BF | EGK.VI-IX | NS | bf_14 | 84 | 6 (7.1%) | 0 | 9 (10.7%) | 2 (2.4%) | |
| BF | EGK.VI-IX | NS | bf_15 | 511 | 51 (9.9%) | 1 | 36 (7.0%) | 4 (0.8%) | |
| BF | EGK.VI-IX | WM | bf_16 | 549 | n/a | n/a | n/a | n/a | 45 |
| BF | EGK.VI-IX | WM | bf_17 | 504 | n/a | n/a | n/a | n/a | 54 |
| BF | EGK.VI-IX | WM | bf_18 | 336 | n/a | n/a | n/a | n/a | 26 |
| BF | EGK.VI-IX | WM | bf_19 | 482 | n/a | n/a | n/a | n/a | 39 |
| ZF | HH4 | TC | zf_15 | 836 | 0 | n/a | 15 (1.8%) | 9 (1.1%) | |
| ZF | HH8 | TC | zf_16 | 608 | n/a | n/a | n/a | n/a | 0 |
| BF | HH4 | NS | bf_20 | 1120 | 27 (2.4%) | 0 | 0 | 3 (0.3%) | |

[1] Zebra finch (ZF), Bengalese finch (BF)

[2] Whole-mount (WM), nuclei spreads (NS), transverse cryosections (TC), longitudinal cryosections (LC)

The numbers of GRC-positive nuclei and micronuclei on cell spreads after FISH with the GRC probe were counted on a randomly chosen area, covering around 50% of the slide. To count GRC-positive nuclei and micronuclei on cryosections we chose 5-6 sections evenly distributed through embryo bodies. For whole-mount, we analyzed cells in ~2 visible (detectable) outer cell layers.

[3] The number of nuclei with two GRC signals may be underestimated due to possible signal overlap.

[4] For nuclei spreads, the number of GRC-positive micronuclei on the slide was counted. For whole-mount embryos and cryosections, the number of GRC-positive micronuclei inside the cell cytoplasm was counted.

* the number of GRC-positive nuclei and micronuclei in EGK.IV zebra finch embryo could not be estimated due to insufficient probe penetration to these cells

** the number of micronuclei was not counted because their morphology was poorly preserved.

**Table S2.** List of antibodies used in this study.

| Species | Antibody | Host | Supplier (catalog #) | Dilution | Reaction type |
|---|---|---|---|---|---|
| Zebra finch, Bengalese finch | Anti-H3K9me3 | Rabbit | Abcam (ab8898) | 2:100 | Unconjugated |
| Zebra finch, Bengalese finch | Anti-Lamin B1 | Rabbit | Abcam (ab16048) | 2:100 | Unconjugated |
| Zebra finch | Anti-BAF | Rabbit | Abcam (ab129184) | 1:100 | Unconjugated |
| Zebra finch | Anti-DAZL | Rabbit | Invitrogen (PA5-54079) | 1:100 | Unconjugated |
| Zebra finch | Anti-rabbit | Donkey | Jackson ImmunoResearch (711-095-152) | 2:100 | FITC |
| Zebra finch | Anti-rabbit | Goat | Jackson ImmunoResearch (111-165-144) | 0.5:100 | Cy3 |
| Zebra finch, Bengalese finch | Anti-H3S10p | Rabbit | GeneTex (GTX128116) | 0.5:100 | Unconjugated |
| Zebra finch, Bengalese finch | Anti-tubulin | Mouse | Abcam (ab7291) | 2:100 | Unconjugated |
| Zebra finch, Bengalese finch | Anti-rabbit | Goat | ThermoFisher Scientific (A-11008) | 0.5:100 | Alexa Fluor™ 488 |
| Zebra finch, Bengalese finch | Anti-mouse | Goat | ThermoFisher Scientific (A-11012) | 0.5:100 | Alexa Fluor™ 594 |

### Supplementary Methods

#### Preparation and testing of *L. domestica* GRC probe

To identify GRC-specific tandem repeat, we isolated DNA from the testis and liver from one male of Bengalese finch using a phenol-chloroform extraction method as described before (Pajer et al. 2006; Schlebusch et al. 2023). The DNA samples were sent to SEQme company (Dobris, Czech Republic) for the 10x Genomics linked-read library preparation and 2x150 bp paired-end sequencing using the NovaSeq 6000 (Illumina). The resulting sequencing reads were trimmed using the longranger basic tool v2.2.2 (10xGenomics) and the testis reads were uploaded to Galaxy server environment for repetitive elements identification using RepeatExplorer2 (Novak et al. 2010, 2013) including the tandem repeats identification using the embedded TAREAN tool (Novak et al. 2017). To identify a GRC-specific repeat, which could be used for the development of GRC probe, we searched for testis repeats that were missing from the liver sequencing reads dataset. Using BWA v v0.7.17 (Li 2013) we mapped the reads from liver on the resulting RepeatExplorer2 contigs combined with the sequences of the TAREAN tandem repeats. Samtools depth v1.14 (Danecek et al. 2021) was used to calculate the coverage along each predicted testis repeats. We identified a 17-bp long tandem repeat (TAAGGACACGAGCTGGG), which was present in 3.8% of the analyzed testis reads and had virtually no coverage (0.24x) in the liver. This made it the best candidate for the GRC-specific FISH probe preparation.

We further used FISH to cytogenetically verify that the candidate repeat is located on GRC. We used two kinds of probes against its sequence. First, we used probes synthesized and labeled by PCR using the following primers (annealing temperature 61°C): Forward 3`-GGACACGAGCTGGGTAAGGACACG-5`, Reverse 3`-CCCAGCTCGTGTCCTTACCCAGC-5`; PRC probes were labeled with biotin-16-dUTP (Roche, Mannheim, Germany) or digoxigenin-11-dUTP (Roche, Mannheim, Germany). Second, we used the following commercially synthesized oligoprobe targeting the GRC-specific repeat was prepared (3`-GGACACGAGCTGGGTAAGGACACG -5`) and labeled with digoxigenin-11-dUTP. To check the specificity of the probes, we tested them on nuclei and synaptonemal complex spreads from male gonads.

Synaptonemal complex spreads were obtained from testes of reproductively active males, following (Peters *et al.*, 1997) with modifications (Poignet *et al.*, 2021). After hypotonic treatment (30 mM Tris, 50 mM sucrose, 17 mM trisodium citrate dihydrate, and 5 mM EDTA; pH 8.2), testes were transferred to 100 mM sucrose and disaggregated. The resulting cell suspension was dropped onto a slide treated with 1% PFA and 0.15% Triton X100 (Sigma Aldrich). After incubation, slides were rinsed in 1× PBS, and immunostained.

Lateral components of synaptonemal complexes were visualised by rabbit polyclonal anti-SYCP3 antibody (ab15093, Abcam, dilution 1:200); centromeres were visualised by human anticentromere serum (15-234, Antibodies Incorporated, dilution 1:50). We used the following secondary antibodies: Alexa-594-conjugated goat anti-Rabbit IgG (H+L) (A32740, Invitrogen; dilution 1:200) and Alexa-488-conjugated goat anti-Human IgG (H+L) (A-11013, Invitrogen; dilution 1:200). Antibodies were diluted in PBT (3% BSA and 0.05% Tween 20 in 1× PBS) and incubated in a humid chamber for 90 min. Slides were then washed in 1× PBS, dehydrated in ethanol (50%, 70%, and 96%, 3 min each), dried and stained with DAPI in mounting medium Vectashield (Vector Laboratories).

Both probes specifically labelled the GRC (*SI Appendix,* Fig. S4). For the analysis of GRC in Bengalese finch embryos, we used oligoprobe labeled with digoxigenin.

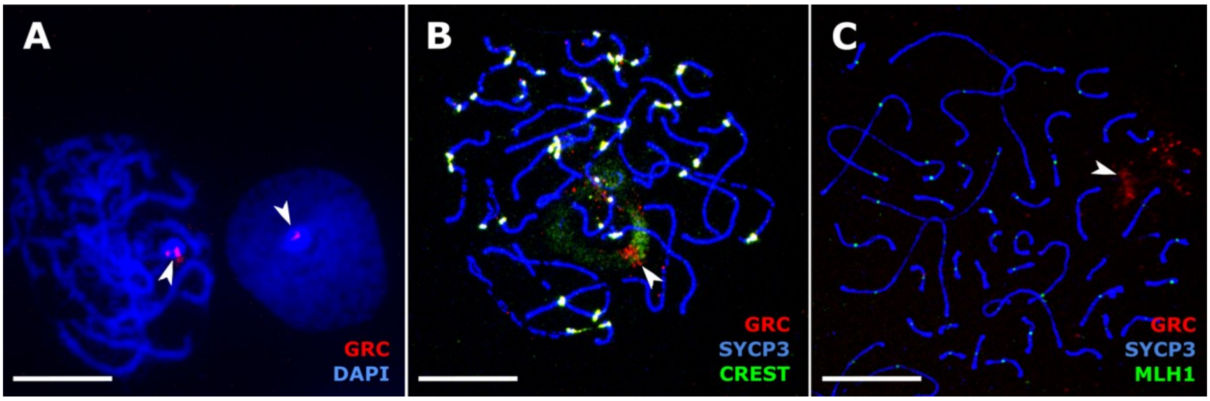


**Fig. S4. FISH-based detection of Bengalese finch GRC on nuclei (A) and synaptonemal complex (B, C) spreads.** GRC-specific repeat (red) is visible as a bright signal on nuclei spreads (A) and as diffuse signals in synaptonemal complexes (B, C). Synaptonemal complexes were visualised using antibodies against SYCP3 protein (blue in B, C); centromeres and GRC were stained by CREST antibodies (green in B); recombination loci were detected by MLH1 antibodies (green in C). Arrowheads point to GRC. Scale – 10 µm.
